# A synoptic solution to a Wright-Fisher model with recurrent mutation, drift and selection

**DOI:** 10.1101/2025.08.27.672576

**Authors:** Philip G. Madgwick, Ricardo Kanitz

## Abstract

The time to fixation (conditional upon fixation) and probability of fixation are key summary statistics of classical population genetics. Here, they are integrated to solve the time to fixation unconditional on fixation in a Wright-Fisher model with recurrent mutation, drift and selection. In its derivation, minor improvements to the probability of fixation and the probability distributions of emergence and spread (or passage) are made. The solution is an exponentially modified Gaussian (ex-Gaussian) distribution with a mean and variance that is the sum of the mean and variance of the underlying exponential and normal (i.e. Gaussian) distributions. To demonstrate its value as a synoptic solution to the Wright-Fisher model, the ex-Gaussian distribution is used to partition the effects of the evolutionary forces of mutation, drift, selection and (to some extent) migration on chosen points of the probability distribution of fixation times. Further, the solution is used to derive (and show the quantitative meaning of) the stochastic drift barrier to nearly neutral mutations with very low selection coefficients, and its consequences on the probability distribution of phenotypic and/or fitness effects that contribute to adaptation. In this way, the probability distribution of the time to fixation unconditional on fixation provides an elegant summary of some key results from classical population genetics, and is also likely to have many other theoretical and applied uses.

## Main-text

The fixation time is a key statistic of classical population genetics ((Crow & Kimura, 1970) p.418), which describes how long it takes for a mutation to spread through a population. Its utility as the go-to summary statistic of classical populations genetics is impeded by an underlying assumption of its derivation, which makes the estimated time to fixation conditional upon fixation (Kimura & Ohta, 1969). Consequently, another key statistic provides a complementary insight, which is the probability of fixation (Kimura, 1962). In a regime of recurrent mutation, both estimates could be described as a single summarisation through the probability distribution of the time to fixation unconditional upon fixation. But, to the best of our knowledge, there is currently no such a summarisation because of the challenges in deriving a closed-form solution (see (Ewens, 2004) especially p. 20-31 and p.92-99). Here, one is provided, followed by a demonstration of its utility as a summarisation.

Consider a Wright-Fisher model of a single locus in a population of constant size that evolves over discrete generations through a mutant allele competing against a wildtype allele under recurrent mutation, drift and selection (Fisher, 1930; Wright, 1931). In each generation, one or more mutations may occur following a binomial process where the mutation rate (*μ*) is the probability of success and the allele number (*cn* where *c* is the ploidy and *n* is the population size) is the sample size. Allele frequency change occurs through the multinomial sampling of genotypes based on their fitness-weighted frequencies, with mutant fitness is described using a selection coefficient (*s* where fitness is 1 + *s*). This model has been the focus of intense study (as summarised in (Ewens, 2004)), including a previous analysis that has shown that it is possible to derive highly accurate approximations of the evolutionary outcomes of this model by converting it into continuous generations to derive closed-form solutions (Madgwick *et al*., 2024). Some methodological improvements we propose here give the way to a more general solution, with special attention given to the case of moderate-to-strong selection.

A statistic that is used throughout the derivation is the probability of fixation. The classic solution (Kimura, 1962) comes from the diffusion approximation (*p* = (1 − *e*^−2*s*^)/(1 − *e*^−2*cns*^)), which (though little discussed) is only accurate under weak selection (*s* < 0.1). A more general solution can be derived, utilising the methods that derive the exact closed-form solution of the fixation probability in a Moran model (Moran, 1958) using a Poisson point (i.e. branching) process (see (Ewens, 2004) p.104-109, and especially eqn. 3.67). A Moran model can be converted into a Wright-Fisher model by rescaling the (drift) variance in allele frequencies per time step, which differs by a factor of two, and permitting allele frequencies to increase by more than one copy per time step, which scales with the strength of selection (*αs*) to a second-order approximation (of the binomial process):

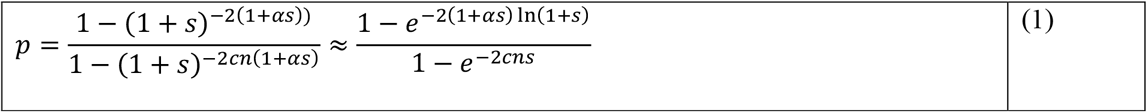

The scalar (*α*) can be statistically fitted for any population size, though it is largely a property of the model (e.g. *α* = {0.146722 − 0147766} for *n* = {10^3^ − 10^6^}; also note the infinite approximation from Taylor series expansion is *α* = 1/6). This solution is highly accurate for any selection coefficient (Figure 1). Note that a wide interpretation can be given to the frequency threshold that is chosen to correspond to fixation in this derivation, equally applying to any non-rare frequency threshold because only rare alleles may be lost by drift (as explained further later).

**Figure 1.**
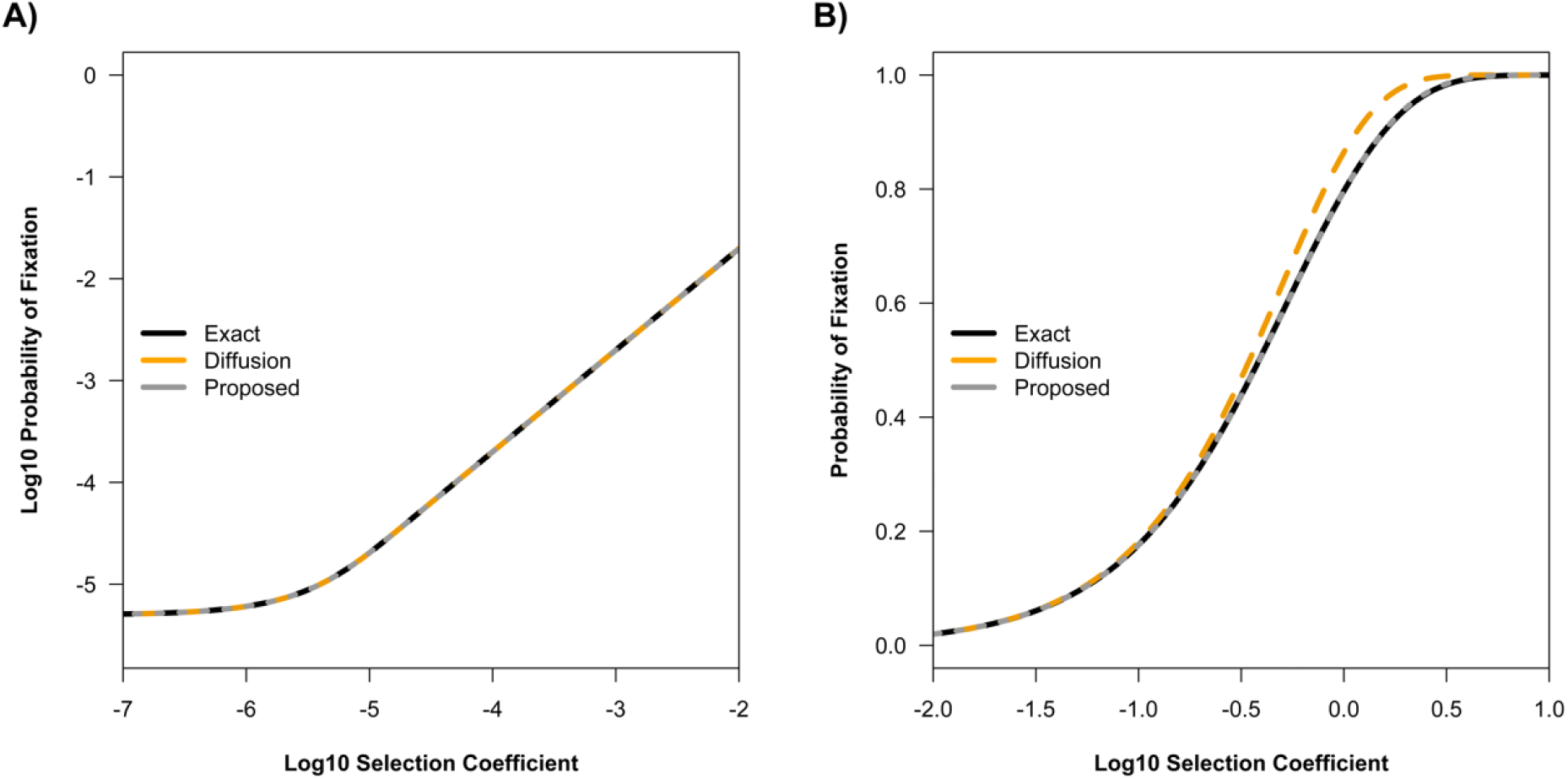
The (log10) probability of fixation from different estimates across log10 selection coefficients, under A) weak selection and B) moderate-to-strong selection for example parameters: *n* = 10^5^, *μ* = 0.1/2*n, s* = 1, *m* = 0, *f*_*T*_ = 1 − (1/2*n*).

The probability distribution of emergence times describes the origin a mutation conditional upon establishment, such that an allele arises and will spread through the population. Whilst the probability distribution of any number of mutations within a generation follows a binomial distribution, the probability distribution of a mutation occurring for the first time in a generation has a geometric distribution, parameterised by the probability and sample size of the underlying binomial process. The probability of one or more mutations (*u*) that will spread is dependent on the product of the independent probabilities of mutation (*μ*) and fixation (*p*) as sampled by the number of alleles in the population (2*n*):

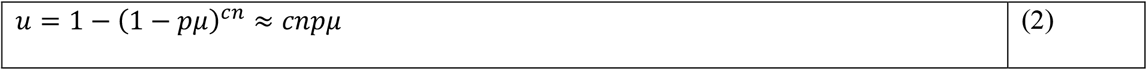

As previously neglected, in the conversion to continuous generations, when the sample size is large by statistical standards (*cn* > 30, which is very small for population sizes), the geometric distribution approaches an exponential distribution:

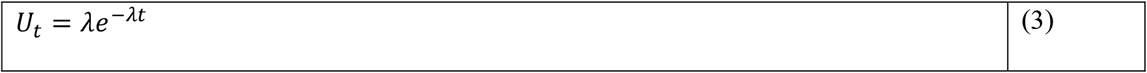

Where *λ* = − ln(1 − *u*), which is also the reciprocal of the mean and standard deviation of the exponential distribution (1/*λ*).

The probability distribution of spreading (or passage) times describes how long it takes a mutation to reach a threshold frequency. As previously neglected, the probability distribution of the time to a frequency threshold logically has a Poisson distribution as a waiting-time process (see also (Ewens, 2004) p.29-30 especially eqn. 1.81), parameterised by a mean (without needing a separate derivation of the variance because it is the same). The mean spreading time can easily be derived from deterministic dynamics (of the logistic increase in allele frequency), which can provide an exact solution under genic selection (see (Crow & Kimura, 1970) p.173-195 especially eqn. 5.3.12). But this derivation assumes that drift has a symmetric effect in increasing or decreasing allele frequencies. To account for the loss of rare alleles, the allele frequency change can be divided by the fixation probability (Kimura & Ohta, 1969):

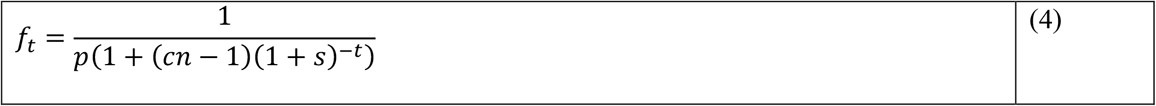

Solving the equation, the mean spreading time to the frequency threshold (*f*_*T*_) conditional upon fixation is:

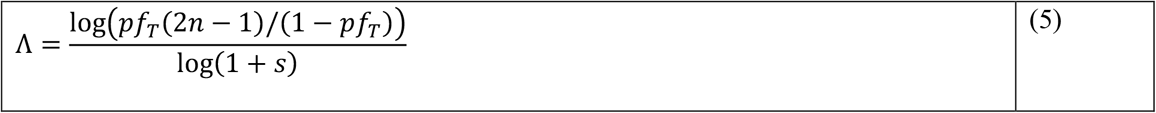

As previously implicit in the conversion to continuous generations, when the mean spreading time is large (Λ > 10), the Poisson distribution approaches a normal distribution (*φ*), following the central limit theorem:

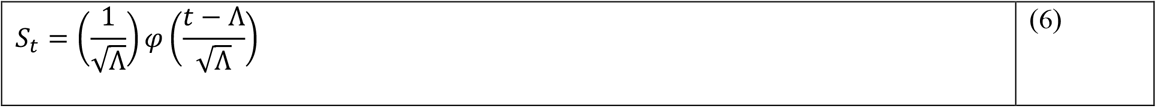

Following its basis as a Poisson distribution, the mean and variance are the same (Λ). Note, again, that the exact frequency threshold (*f*_*T*_) that is chosen is of little importance to the derivation. Here, it can be clearer why, because the coefficient of variation 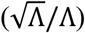 drastically decreases as the frequency threshold of the allele copy number increases.

Previously, at this juncture, analytical techniques have had to give way to numerical techniques to describe the convolution of the emergence and spreading time distributions (Madgwick *et al*., 2024).

However, as has been neglected in population genetics to the best of our knowledge, the sum of random variables sampled from an exponential and normal (i.e. Gaussian) distribution have an exponentially modified Gaussian (ex-Gaussian) distribution (Hohle, 1965), which is well-characterised (using *ϕ* as a cumulative normal distribution):

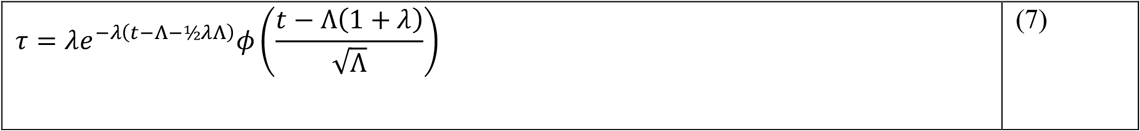

The ex-Gaussian distribution is obscure, having been derived by cognitive psychologists to study reaction times – and scarcely applied outside of this context. The solution describes the probability distribution of the time to a frequency threshold unconditional on reaching that frequency threshold; the integral has previously been proposed but not solved (e.g. (Ohta & Kimura, 1971)). The ex-Gaussian distribution is a continuous generations equivalent of the geometric-Poisson (or Pólya–Aeppli) distribution, which has also been implied (e.g. (Ewens, 2004) eqn. 2.146) but lacks a closed-form solution (meaning that, e.g., the mean and variance cannot be directly inferred). By contrast, the mean and variance of the ex-Gaussian distribution is the sum of the means and variances of the component distributions ((1/*λ*) + Λ; (1/*λ*^2^) + Λ), often having the general shape of a skewed normal distribution with a long tail (Figure 2).

**Figure 2.**
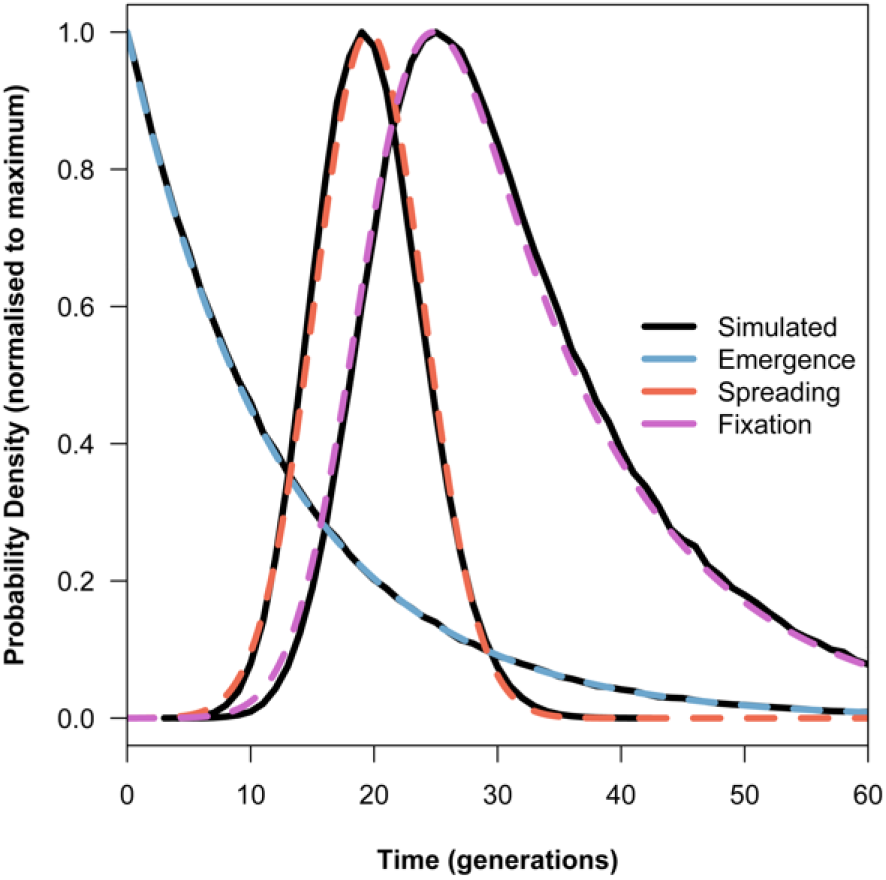
The probability distribution of the time to emergence, spreading and fixation for example parameters: *n* = 10^5^, *μ* = 0.1/2*n, s* = 1, *m* = 0, *f*_*T*_ = 1 − (1/2*n*).

Why is this useful as a synoptic solution to the Wright-Fisher model? The equation neatly encapsulates the net effects of mutation, drift and selection on the spread of a mutant allele against a wildtype allele at a single locus. The probability distribution of the time to fixation unconditional on fixation brings together the key statistics of classical population genetics into a single summarisation. Consequently, it is possible to go further than previous analysis in quantifying the effects of mutation, drift and selection on the fixation time. Moreover, this analysis can be undertaken for different points on the probability distribution to quantify the effects under different stochastic outcomes. For the purposes of demonstration, take the frequency threshold as fixation (*f*_*T*_ = 1 − 1/*cn*), assume genic selection (i.e. the homozygote mutant has fitness (1 + *s*)^2^) and consider the following chosen points of the probability distribution: the minimum at 2.5% cumulative probability density, the mode, the median, the mean and the maximum at 97.5% cumulative probability density. Note that it is useful that the exponential, normal and ex-Gaussian distributions all have well-characterised cumulative functions for precisely computing the chosen points. The ex-Gaussian cumulative density function is:

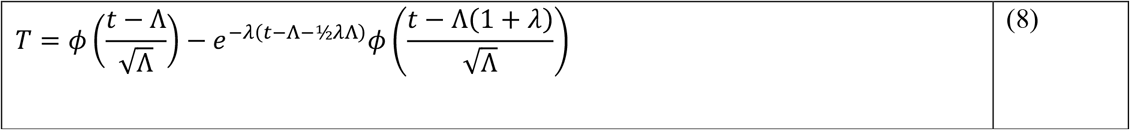

Like the probability distribution, the solution is exact.

The fixation time can be partitioned into effects from mutation, drift and selection. Drift has opposing effects on emergence and spreading (see eqn. 2 and 4) that are worth separating: slowing down the emergence time through the chance loss of rare alleles and speeding up the spreading time by increasing allele frequency conditional upon fixation. The roles of selection and mutation can be separated from drift by removing the fixation probability from the calculation of the fixation time (by setting *p* = 1 in eqn. 3 and 6). With the removal of drift, the probability distribution of the fixation time is the probability distribution of the emergence time displaced by the spreading time, which (by solving for *t* = *T* in eqn. 4 when *f*_*t*_ = *f*_*T*_) has an invariant (i.e. deterministic) contribution to any chosen point of the probability distribution of fixation times. Once drift has been removed from the emergence time, the cumulative probability distribution for the emergence times (1 − *e*^−*λt*^) can be used to calculate the mean contribution of mutation to the chosen points of the probability distribution for the time to fixation unconditional on fixation. To describe the role of drift for any fixation time, it is possible to work backwards to infer the probability distribution of the underlying emergence and spreading times by calculating the probability of sampling a chosen value to make the emergence and spreading times sum to the chosen fixation time (Figure 3). The mean contributions of drift to emergence and spreading times can then be calculated as the difference between the means of the underlying emergence and spreading times and the means from mutation and selection. Thereby, the fixation time for different chosen points is partitioned into mean contributions from mutation, drift (during emergence and spreading) and selection (Figure 4A). Interestingly, despite having opposing effects on the emergence and spreading times, the dominant effect of drift varies with the strength of selection; under stronger selection, drift is more likely speed up the fixation time, as it does for all chosen points in the example (where *s* = 1).

**Figure 3.**
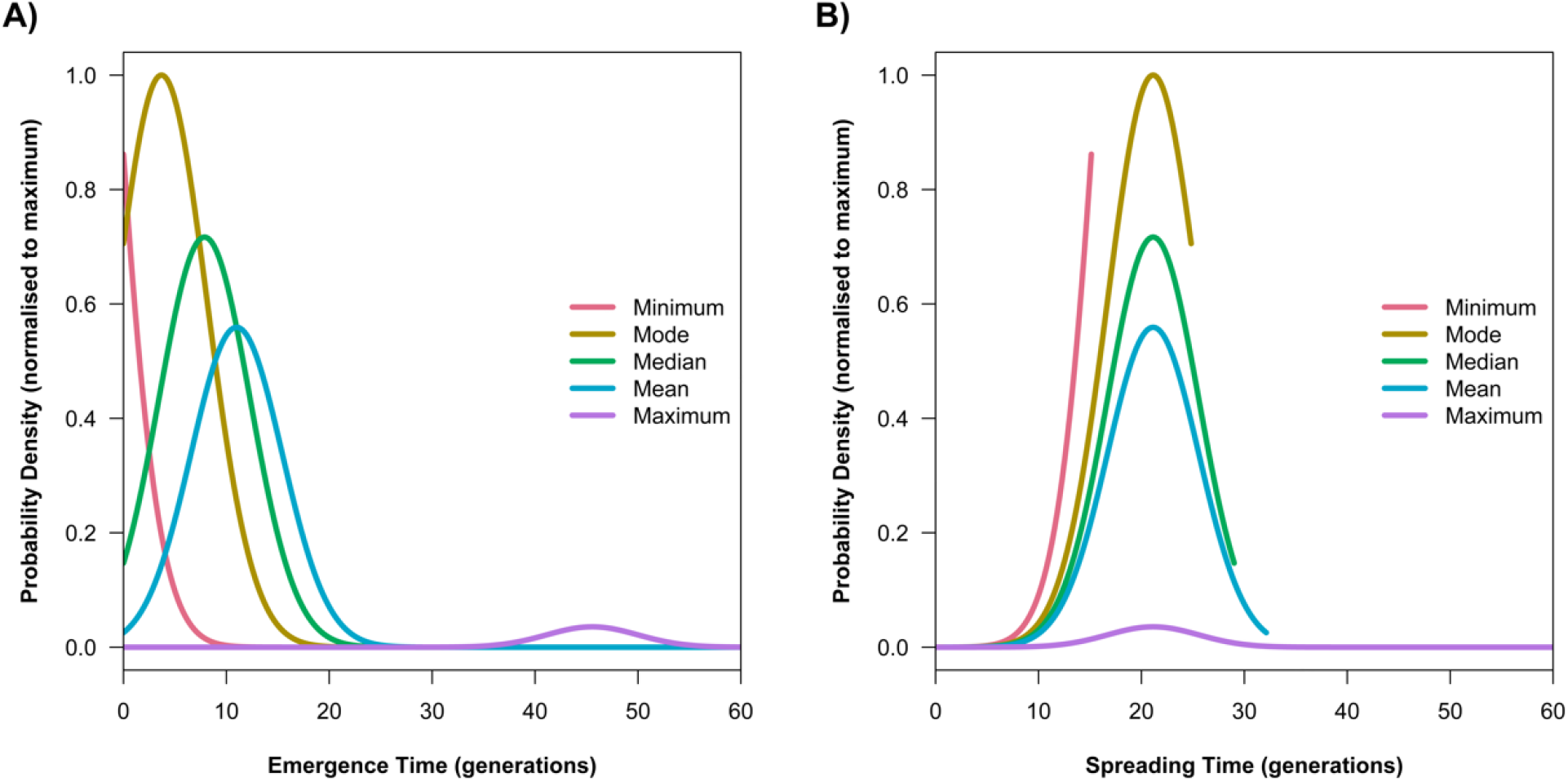
The probability distributions of A) emergence and B) spreading times that can give rise to a particular times to fixation (namely: the minimum at 2.5% cumulative probability density, the mode, the median, the mean and the maximum at 97.5% cumulative probability density; which also define the truncations) for example parameters: *n* = 10^5^, *μ* = 0.1/2*n, s* = 1, *m* = 0, *f*_*T*_ = 1 − (1/2*n*).

**Figure 4.**
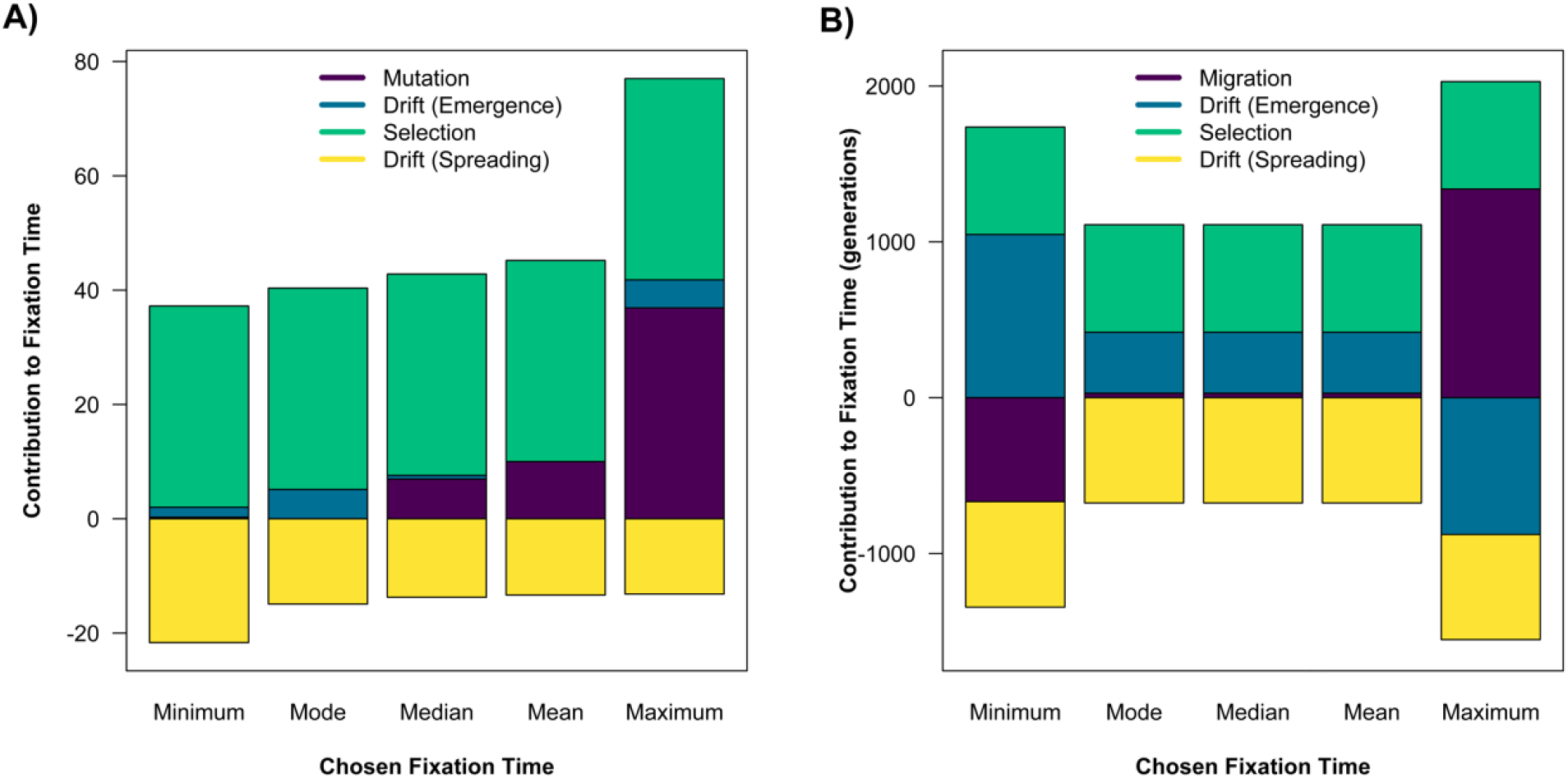
The contributions of evolutionary forces to some chosen points of the probability distribution of the time to fixation (namely: the minimum at 2.5% cumulative probability density, the mode, the median, the mean and the maximum at 97.5% cumulative probability density; and, note, the total fixation time is the highest positive bar minus the lowest negative bar), with emergence by: A) mutation for example parameters *n* = 10^5^, *μ* = 0.1/2*n, s* = 1, *m* = 0, *f*_*T*_ = 1 − (1/2*n*) or B) migration for example parameters *n* = 10^3^, *z* = 30, *μ* = 0.01/2*n, s* = 1, *m* = 10^−2^, *f*_*T*_ = 1 − (1/2*n*) (and note probability of emergence by migration (not mutation) for these parameters is: *e*^−*λV*^ ≈ 0.89; see eqn. 12).

To complete the analysis of the basic evolutionary forces identified by classical population genetics using the solution, it is possible to extend the basic setup of the Wright-Fisher model to include migration using a simple stepping-stone model (Kimura & Weiss, 1964). Suppose that a distal population has a copy of a beneficial allele, but there are *z* − 1 stepping-stone populations (for *z* ≥ 1) separating it from a focal population. The probability distribution of the allele arising by mutation in the focal population has already been described (eqn. 3), albeit that now 1 + *s* must be substituted with (1 + *s*)(1 − *m*) using the migration rate *m* to account for emigration out of the focal population. The probability distribution of the allele spreading from one population to another by migration can be derived in a similar way to the probability of mutation (*u*; eqn. 2) using the probability of migration (*θ*), giving a probability of one or more migrant individuals in the population (*n*, not 2*n* as before for the allele population size) moving towards the focal population (at rate ½ because of bi-directional migration) of:

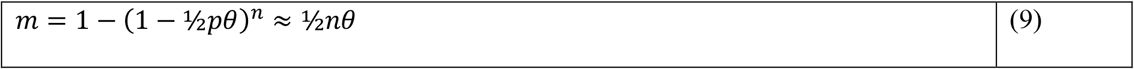

The geometric distribution of the time to first-migration (or invasion) approaches an exponential distribution when the sample size is large by statistical standards (*n* > 30):

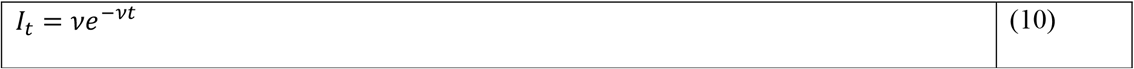

Where *ν* = − ln(1 − *m*), which is also the reciprocal of the mean and standard deviation of the exponential distribution (1/*ν*). It is necessary, at this juncture, to assume that the allele does not migrate between populations when it is rare, and rapidly transitions between a negligible frequency and near-fixation (as if *f*_*T*_ > ½ is *f*_*T*_ ≈ 1) following the probability distribution that has also previously been described (eqn. 6; as if under strong selection). The sum of *z* random variables sampled from this exponential distribution is a gamma distribution (with shape *z* and rate *ν*), which approaches a normal distribution (with mean *z*/*ν* and variance *z*/*ν*^2^) for a statistically large number of samples (*z* > 10). Altogether, the probability distribution of the time to the allele arising by migration in the focal population is:

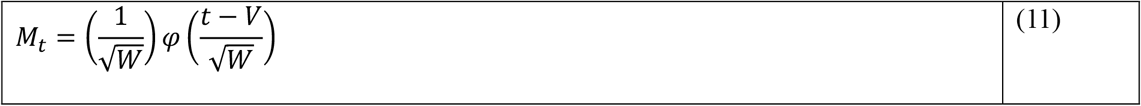

Using mean *V* = *z*((1/*ν*) + Λ) and variance *W* = *z*((1/*ν*^2^) + Λ). The cumulative probability distribution over *z* describes the stochastic wave of advance. The probability distribution of the time difference between random variables sampled from the probability distribution of migration and mutation can then be calculated, which has an ex-Gaussian distribution. The overall probability of the allele arising by mutation (rather than migration) is the sum of all positive values (i.e. *U*_*t*_ − *M*_*t*_ > 0), which can be calculated using the cumulative density function:

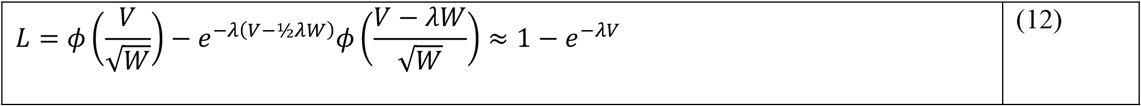

The combined probability of emergence by mutation in the focal population and migration from the distal population is:

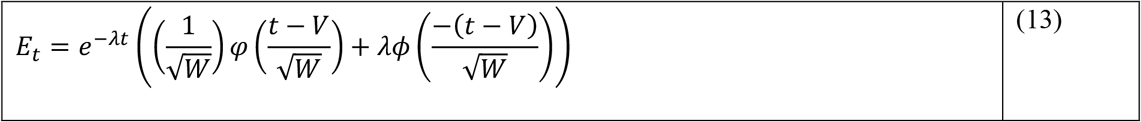

The composite distribution is intractable. Nonetheless, the derivation of the overall probability of the allele arising by mutation (not migration; eqn. 12) provides an indication of the relative importance of migration to the emergence time, and the combined probability of emergence by mutation and migration (eqn. 13) can be directly compared to the emergence time by mutation alone. In many cases, migration would either be important (*L* ≈ 0) because the distal population is sufficiently nearby or not (*L* ≈ 1) because it is sufficiently far away (and there is no exact solution for intermediate cases). When migration is unimportant, the results of the partial effects of evolutionary forces remain the same as before, whereas when migration is important, migration can replace mutation in driving the emergence of the allele. As the sum of normal random variables follows a normal distribution (adding together their means and variances), the time to fixation unconditional on fixation is:

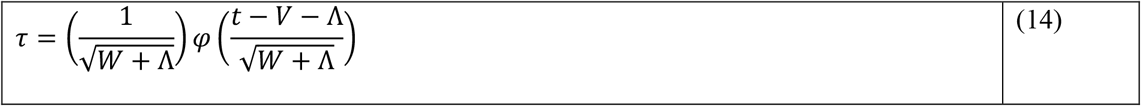

As before, the fixation time for different chosen points can then be partitioned into mean contributions from migration, drift (during emergence and spreading) and selection (Figure 4B), noting that the mode, median and mean are the same for a normal distribution. In comparison to the previous analysis, unlike mutation (and more like drift), migration has two opposing effects on the fixation time: it may decrease the effective selection coefficient in the focal population with consequences for drift and selection, which slows down the fixation time, and it may increase the effective mutation rate when the allele arises by migration, which speeds up the fixation time.

Besides changing the fixation time, another important insight is embedded in the probability distribution of the time to fixation unconditional on fixation, which is the drift barrier that makes mutations with very low selection coefficients have a probability and time to fixation equivalent to a neutral mutation (Kimura & Ohta, 1969). The ex-Gaussian distribution can be used to derive the drift barrier because it may fail to be computed when the mean of the underlying normal distribution is undefined (eqn. 5) because the argument of the numerator (i.e. inside the log) is less than one (when using the 2*s* approximation of the fixation probability for small *s*). Solving for the selection coefficient, the drift barrier for selection coefficient (*s*\*) can be approximated as:

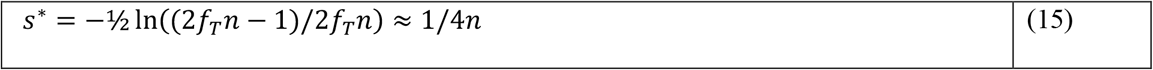

The result is slightly lower than the previous derivation of 1/2*n*, which also uses the 2*s* approximation (Kimura, 1968; Kimura & Ohta, 1969). The difference is of little importance when it is recognised that both estimates are placing a threshold on a continuum when using the full solution for the fixation probability (Figure 5).

**Figure 5.**
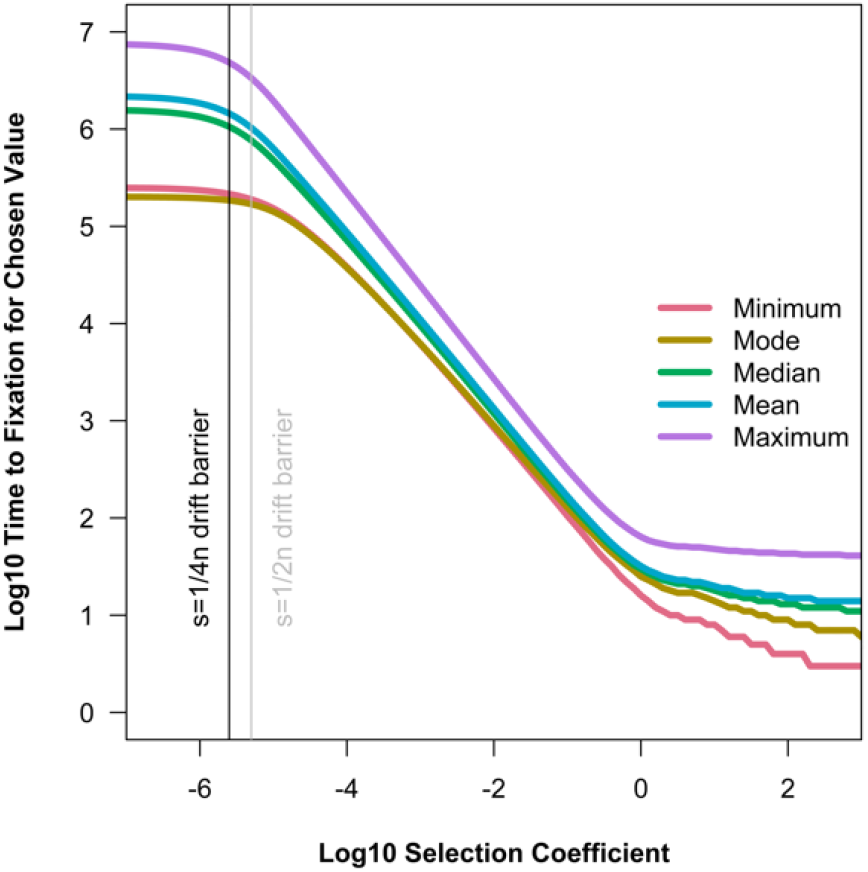
The time to some chosen points of the probability distribution of the time to fixation unconditional on fixation (namely: the minimum at 2.5% cumulative probability density, the mode, the median, the mean and the maximum at 97.5% cumulative probability density) across selection coefficients given in log10 units for example parameters: *n* = 10^5^, *μ* = 0.1/2*n, s* = 1, *m* = 0, *f*_*T*_ = 1 − (1/2*n*) (and note the black and grey lines show the predicted drift barriers; see eqn. 15).

The effect of the drift barrier has been influentially presented in another way, reflecting the probability distribution of phenotypic and/or fitness effects that contribute to adaptation. In the geometric model of adaptation, the probability that a mutation is favourable decreases as its standardised effect size (*x*) increases (Fisher, 1930). When the number of phenotypic dimensions is high (> 5), the probability of a mutation being favourable approaches the infinite solution (*ϕ*(−*x*)), which would suggest that smaller mutations are more important to adaptation. However, when factoring in the fixation probability (*p*) assuming that phenotypic effects are proportional to fitness effects (*pϕ*(−*x*)), intermediate mutations are more important to adaptation intermediate mutations are more important to adaptation (Kimura, 1983). Using the more accurate fixation probability (eqn. 1) and cumulative ex-Gaussian distribution (eqn. 8), it is possible to be more precise, by solving the probability of a mutation having fixed within a timeframe (*Tϕ*(−*x*)). Note that when the timeframe is infinite, the solution simplifies (to (1 − *e*^−*λ*(*t*−Λ−½*λ*Λ)^)*ϕ*(−*x*)). Rather than considering sequential fixations in a regime of strong selection and weak mutation (like (Orr, 1998)), which is alien to the philosophical foundations of the geometric model (Fisher, 1930), consider the simultaneous evolution of new mutations at loci with independent selection coefficients. For an infinite number of loci, the expected rate of fixations across phenotypic effects is proportional to:

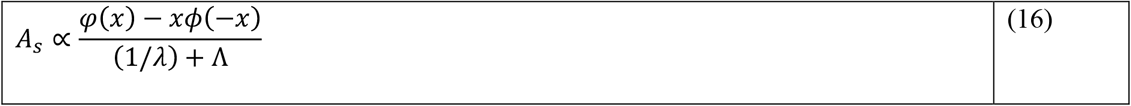

Where *φ*(*x*) − *xϕ*(−*x*) is the contribution of the mutation to adaptation (taking into account *ϕ*(−*x*) as the probability of being favourable; (Orr, 1998)) and (1/*λ*) + Λ is the expected time to fixation (unconditional on fixation). Taking a specific relation between phenotypic and fitness effects (*x* = *s*, following (Kimura, 1983)), it is possible to renormalise this adaptive rate as a probability distribution of mutations of different sizes contributing towards adaptation (Figure 6). Comparing solutions, the adaptive rate suggests that larger mutations can have a greater role in adaptation (because, when they are favourable, they tend to reach fixation faster).

**Figure 6.**
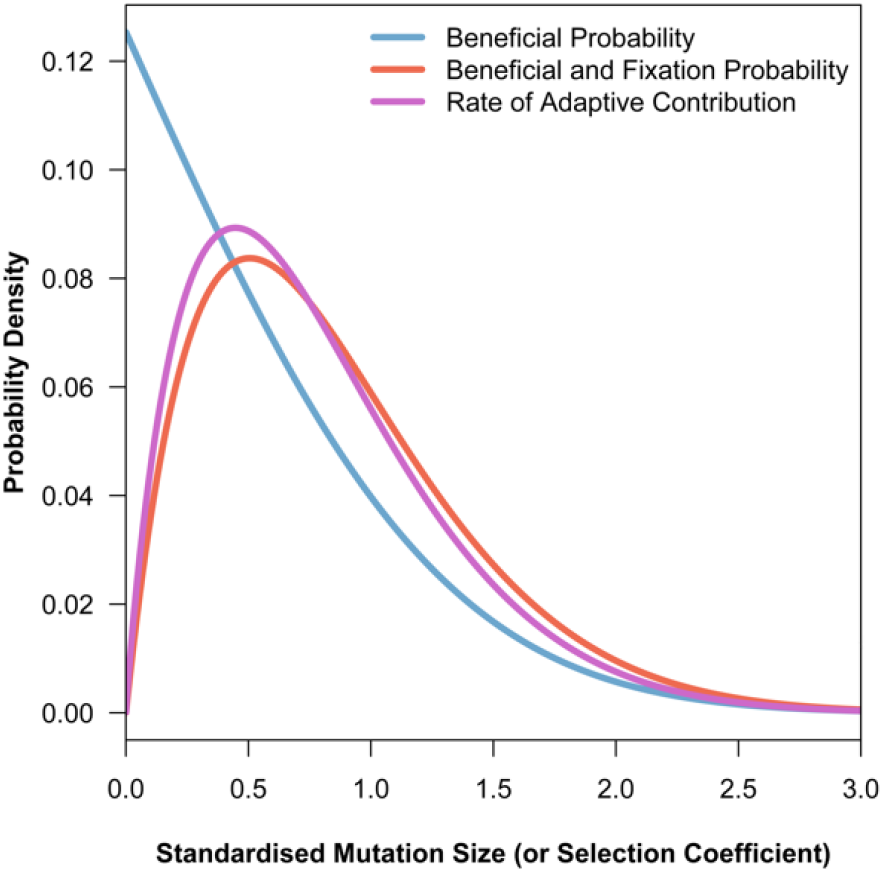
The probability distribution of phenotypic or fitness effects contributing towards adaptation, assuming that the standardised mutation size is equal to the selection coefficient (*s*), for three applications of the geometric model of adaptation: the probability of a mutation being beneficial in the *ϕ*(−*s*), the probability of a mutation being beneficial and reaching fixation ((1 − *e*^−2*s*^)*ϕ*(−*s*)), and the rate that mutations contribute to adaptation for the example parameters *n* = 10^5^, *μ* = 0.1/2*n, m* = 0, *f*_*T*_ = 1 − (1/2*n*) (see eqn. 16, renormalised as a probability distribution).

Therefore, as neatly described by the summary distribution of the time to fixation unconditional on fixation, the roles of mutation, drift, selection and (to some extent) migration can be quantified using the ex-Gaussian distribution solution to show the effects of the evolutionary forces on dynamics and outcomes in the Wright-Fisher model. The solution provides an elegant summary of some key results of classical population genetics, blending together the insights from the time to fixation conditional on fixation and the probability of fixation. There is, certainly, much more that can be done with the solution, which can be readily employed to address theoretical and applied problems.

## Notes

### Competing Interest Statement

The authors have declared no competing interest.

